# Cell-substrate friction controls biofilm development

**DOI:** 10.1101/2025.07.11.664457

**Authors:** Aawaz R. Pokhrel, Raymond Copeland, Maryam Hejri, Tom E. R. Belpaire, Gabi Steinbach, Siu Lung Ng, Brian K. Hammer, Peter J. Yunker

## Abstract

Bacteria often live in biofilms, surface-attached communities that can form on nearly any surface, from coarse sands to smooth glass. It is often hypothesized that cell-substrate friction can impact biofilm growth and development on these disparate surfaces. However, the experimental difficulty in measuring the friction between a biofilm and its surface has limited our understanding of how friction forces impact emergent colony-level behaviors and morphologies. Here, we demonstrate that increasing cell-substrate friction increases the biofilm contact angle, which, in turn, decreases the horizontal biofilm range expansion rate. We first used individual-based simulations to isolate the impact of friction. In this simple model, we found that increasing friction increases biofilm contact angle, and, in turn, that the contact angle emerges from a simple dynamic force balance between cell-substrate friction, cell-cell steric forces, and cell-cell adhesion, reminiscent of Young’s equation for sessile liquid drops. We developed an approach to directly measure the friction between bacterial colonies and agar surfaces, and found that the friction between the substrate and the biofilm is higher on substrates made with higher agar weight percentage. Additionally, we observed that biofilm contact angles increase with friction, following a Young-equation-like balance. Finally, we found that contact angle increases with agar percentage for a wide range of bacteria, suggesting that the biophysical impact of friction may play a role for a wide array of microbes.

## Introduction

Biofilms are surface-attached communities of microbes, including bacteria, archaea, and fungi, that are found in nearly every natural and built environment on Earth. They can form on diverse surfaces, from natural environments such as river beds, coarse sands, and human skin, to industrial substrates such as pipes, pumps, and filters, to laboratory settings like agar gels (1–3). These surfaces have highly varied physical properties, which have been shown to substantially affect biofilm growth and development (4–8). In the lab, different surface properties are often studied by varying the weight percentage of agar gels, which is expected to affect substrate-colony friction, and is known to cause changes in biofilm morphology and growth (5, 9–11). However, our understanding of the role of friction in biofilm development is limited by the fact that direct measurements of friction between a colony and an agar surface have proven difficult. One of the challenges in measuring such forces is the scale at which they occur. Atomic Force Microscopy (AFM) can measure microscopic forces, such as friction between a single cell and an AFM cantilever tip (12); however, measuring mesoscale forces on soft surfaces, like the friction force experienced as a colony grows and slides over an agar surface, is challenging (13–15). Hence, a thorough understanding of the connection between friction and colony growth remains elusive.

Specifically, the friction between the substrate and the colony may affect the balance between horizontal and vertical growth. The interplay between vertical and horizontal growth was previously shown to play an important role in determining biofilm growth rates (16). This trade-off is physically manifested in the contact angle of the biofilm, the angle it makes with respect to the substrate on which it grows (in analogy to a liquid drop (17, 18)); in fact, the biofilm range expansion rate depends more strongly on the contact angle of the biofilm than the cellular growth rate (16). Crucially, higher cell-substrate friction could lead to larger contact angles as cells accumulate more at the edge (5, 19), which would, in turn, lead to lower horizontal biofilm growth rates. Thus, understanding what factors determine the contact angle of a biofilm is crucial to understand biofilm development.

Here, we investigate the role of colony-substrate friction in determining biofilm contact angles. We begin by using agent-based simulations to analyze how range expansion rates depend on cell-substrate friction, cell-cell adhesion, and contact angle. We found that increased friction and cell–cell adhesion both lead to higher biofilm contact angles, thereby reducing the range expansion rate. Our simulations further demonstrate that this relationship is consistent with a force balance reminiscent of Young’s equation, wherein the contact angle mediates the balance between cell-substrate friction, cell-cell steric forces, and cell–cell adhesion. Next, we test if friction has a similar impact on the biofilm contact angle in experiments. We developed a method to directly measure the friction between a colony and its substrate, and observe that colony–surface friction increases as agar concentration rises. We next measured colony contact angles and found that con-tact angle increases with higher colony-surface friction. We show that the Young’s equation-like force balance holds experimentally, too. Collectively, these findings demonstrate that the colony contact angle increases with friction, suggesting that a dynamic force balance critically influences the range expansion rate of biofilms.

## Results

### Biophysical simulations suggest a dynamic force balance with friction

To determine what biophysical factors control the biofilm contact angle, we performed individual based simulations of reproducing bacteria on a surface. Each bacterium is represented as a circle that can grow and divide (see Fig. 1A). Bacteria interact with each other and with the surface on which they sit via physical forces including: cell–cell steric repulsion and adhesion (*S* and *A*, respectively), cell–substrate friction (*F*), and cell-substrate adhesion (20) (see Methods for more details). While this minimal model does not aim to replicate the full complexity of real bacterial behavior, it allows us to isolate the effects of each physical force on biofilm growth and structure. Despite its simplicity, the model captures essential features of biofilm development, such as the spherical cap morphology (Fig. 1A) (16) and the linear relationship between colony diameter and time (Fig. 1B). To facilitate comparisons, we will characterize simulations by *F*^*^ and *A*^*^, which are the maximum friction and cell-cell adhesion forces, respectively, scaled by the average steric force cells experience at the jamming point (21) (which is the same value for all simulations; see Methods for more details).

**Fig. 1.**
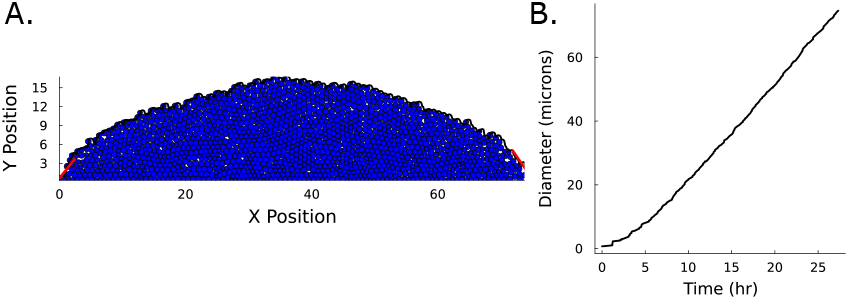
A. A visual representation of our agent-based model with the measurement of the contact angle shown with red lines on either side. B. The diameter of a simulated colony is plotted over a twenty hour time period. We see that after a brief exponential phase the diameter grows linearly over time. This simulation used moderate friction and cell-cell adhesion (*A*^*^ and *F* ^*^ equal to 0.21).

We find that the substrate-cell friction and cell-cell adhesion both play important roles in setting the contact angle, *θ. θ* increases with increasing *F* or *A* (Fig. 2A and inset). In a related vein, increasing *F* or increasing *A* decreases the range expansion rate, *da/dt*, where *a* is the colony radius and *t* is time (Fig. 2B). This result makes intuitive sense; as friction or adhesion increases, cells must grow more before cell-cell steric forces are enough to overcome the forces that resist expansion (i.e., *F* and *A*). If cells cannot be pushed outwards, they are pushed upwards, thus increasing *θ* and decreasing *da/dt*.

**Fig. 2.**
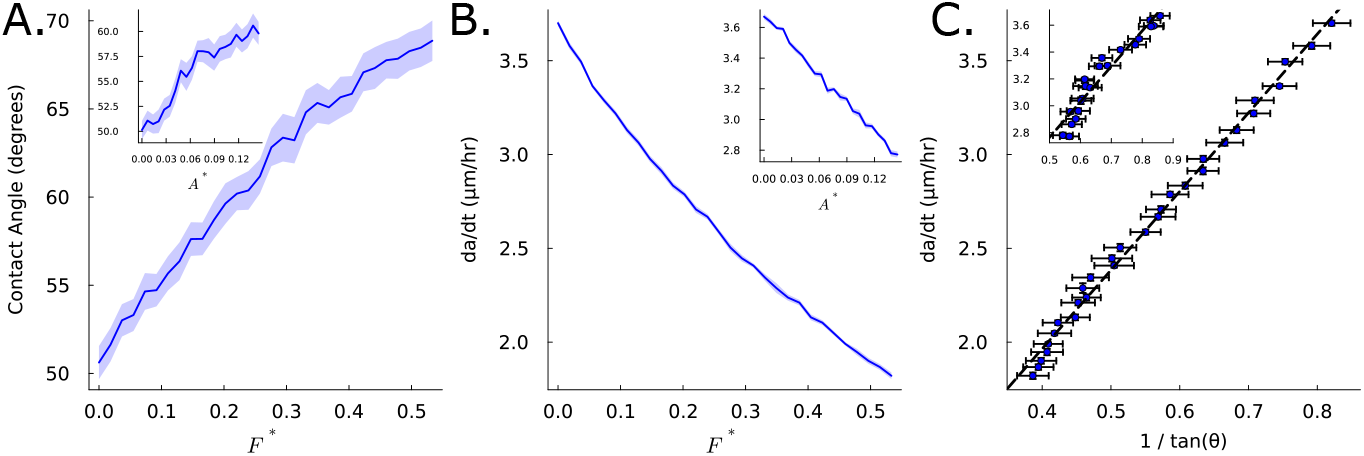
A. Contact angle is plotted as a function of the force of friction (error bars represent standard error n=30). *F* ^*^ is the maximum friction force divided by the average cell-cell steric force at the jamming point where growth rates decrease (see Methods for more details). Inset: Contact angle is plotted as a function of cell-cell adhesive forces. *A*^*^ is the maximum cell-cell adhesion force divided by the average cell-cell steric force at the jamming point where growth rates decrease. B. The mean range expansion rate, *da/dt*, where *a* is the radius and *t* is time, is plotted versus *F* ^*^. Inset: *da/dt* is plotted against *A*^*^. C. We then plot *da/dt* against 1*/* tan(Θ) and find a strong linear relationship as predicted from (16). The linear fit was found without weighting (p value for slope=2.1 · 10^−32^ and *R*^2^ = 0.99). Inset: *da/dt* is plotted as a function of *A*^*^ (p value for slope= 2.2 · 10^−12^ and *R*^2^= 0.92).

Further, these results are consistent with the idea that the edges of colonies can be accurately described as spherical cap napkin rings with constant contact angles (16). In particular, we see that *da/dt* is proportional to 1*/tan*(*θ*), as previously predicted (Fig. 2C) (16).

The spherical cap shape of colonies along with their well-defined contact angles, parallels that of a liquid droplet; therefore, we postulate that a force balance analogous to Young’s equation may govern the contact angle in biofilms (see Fig. 3A for a schematic). In the case of liquid droplets, the contact angle is determined by the balance of interfacial tensions among the substrate, liquid, and air. Drawing upon this analogy, we hypothesize that the cells at the edge of the colonies reach dynamic equilibrium through a similar force balance involving colony-substrate friction, repulsive cell-cell steric forces, and attractive cell-cell adhesive interactions. Specifically, the cell-cell steric forces generated by growth push cells outward, in the direction of colony expansion, while the cell-substrate friction forces directly oppose this motion. Cell–cell adhesion forces also oppose expansion, acting on the cell at the leading edge, on average, at an angle proportional to *θ*.

**Fig. 3.**
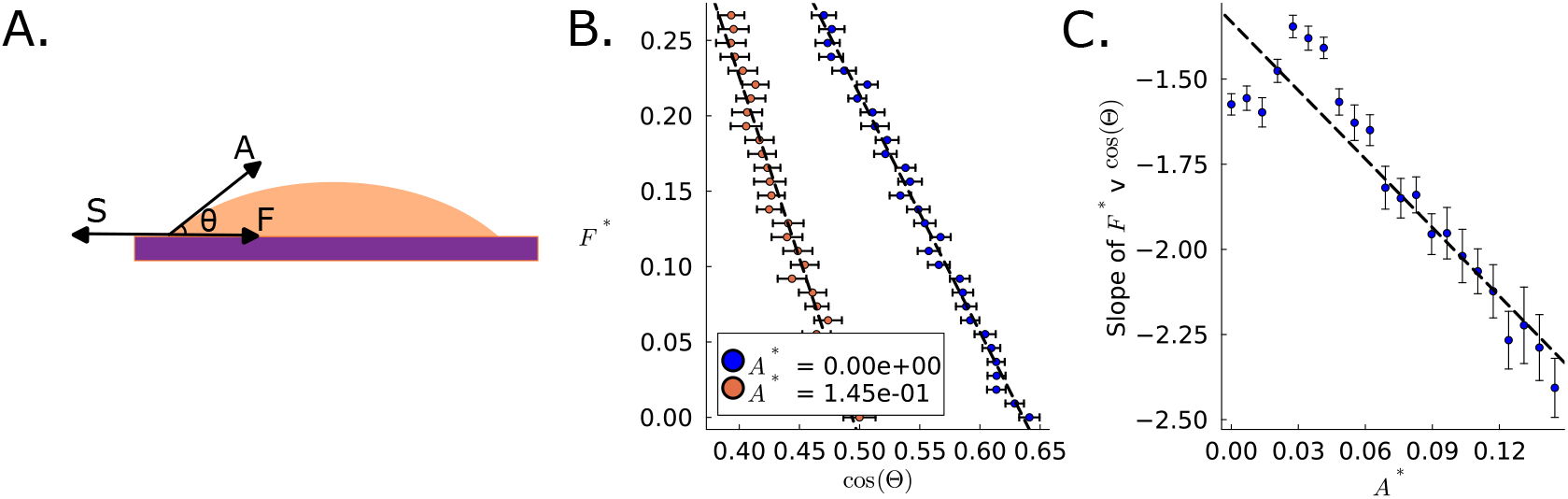
A. Schematic demonstrating the force balance on a cell at the edge of the biofilm between cell-cell steric forces, cell-cell adhesive forces, and cell-substrate friction. B. *F* ^*^ is plotted against *cos*(*θ*). We observe a linear relationship *F* ^*^ and *cos*(*θ*) for the range of cell-cell attraction tested in our simulations (error bars represent standard error, n=40). The best fit lines were unweighted when fit with the means of each point (*A*^*^ = 0 p value for slope=4.9 · 10^−29^ *R*^2^ = 0.98, for *A*^*^ = 5.79 · 10^−1^ p value for slope=6.0 · 10^−22^ *R*^2^ = 0.96). C. We then plot the slope of friction v cos Θ against *A*^*^. The error bars represent the standard error in slope calculation, and the linear fit shown was done with the mean slopes (as shown in B), unweighted (p value for shown line’s slope is 1.8 · 10^−10^ and *R*^2^ = 0.87).

Thus, we propose a simple force balance model inspired by Young’s Equation, where the contact angle is set by balancing friction, steric, and adhesive forces as the colony expands horizontally (see Fig. 3A). This Young’s Equation-like force balance suggests that for a biofilm expanding at a constant rate with a constant contact angle,

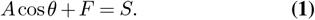

However, biofilms are not equilibrium liquids; buckling instabilities can lead to non-zero contact angles even when *A*^*^ = 0 (Fig. 3A)(22–25). However, if this force balance remains relevant, we would expect that *F* ^*^ should be linear in *cos*(*θ*), with a slope that is negative and proportional to *A*. Thus, to test this idea, we first plotted *F* ^*^ against *cos*(*θ*) (see Fig. 3B). We found that for each value of *A*^*^, *cos*(*θ*) is linear in *F* ^*^ (*R*^2^ *>* 0.9 for all values of *A*^*^). Next, we found that the slope of *F* ^*^ versus *cos*(*θ*) is linear in *A*^*^ (*R*^2^= 0.87), consistent with a Young’s equation-like dynamic force balance. Note, small values of *A* deviate from this linear regime, likely indicating that for them, the buckling of cells impacts the morphology more than cell-cell adhesion.

We next performed sensitivity analysis to determine if *θ* is more sensitive to changes in *F* ^*^ or *A*^*^ (26). The sensitivity of *θ* with respect to *F* ^*^ and *A*^*^ is given by:

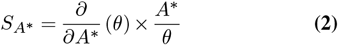

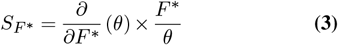

We calculated the ratio of two of these sensitivities,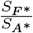,for all of the values of *F* ^*^ and *A*^*^ that we simulated. We chose the range of values for *F* ^*^ and *A*^*^ based on when the forces caused unrealistic biofilm morphology (e.g., *θ >* 90°). We found that 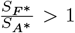 for 72% of the values we studied. In particular, when friction is greater than adhesion (*F* ^*^ *> A*^*^), contact angle always depends more sensitively on friction than adhesion. When friction is less than adhesion (*F* ^*^ *< A*^*^), contact angle sometimes depends more sensitively on friction and sometimes depends more sensitively on adhesion.

### Biofilm-substrate friction increases with increasing agar percentage in experiments

We next sought to experimentally measure the friction force exerted on biofilms as they slide across agar surfaces with various agar percentages. To do so, we developed an experimental method using a Universal Testing Machine (Zwick/Roell) equipped with an X-ForceHP 5N load cell which allowed us to measure forces over a range from 0.05 N to 5 N. We inoculated 4 µl of an OD1 suspension of cells (*Vibrio cholerae* Wild Type, WT) onto a ME Whatman grid filter paper, resulting in a roughly square-shaped deposit, which we then incubated at 23° C for 48 hours. We attached the filter paper to a nylon string with two-part epoxy, and placed the filter paper biofilm side down on an agar pad. The UTM then pulled the nylon string at a constant speed of 5 mm/min, and measured the force necessary to do so (schematic shown in Fig. 4A). We performed these experiments on six different agar percentages (1.5%, 2.5%, 3%, 3.5%, 4.5%, 5%), and performed three replicate experiments for each agar percentage. Note, the friction of an agar pad will depend on how wet or dry the pad is. For these experiments, agar pads were poured at the same time and allowed to dry under the same conditions for the same amount of time.

**Fig. 4.**
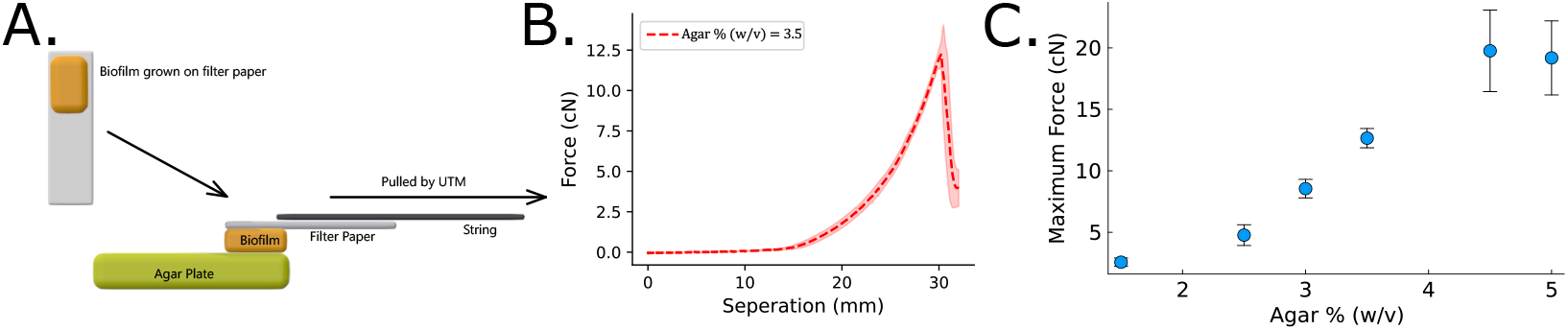
A. A schematic representation of the experimental setup designed for measuring the surface resistance experienced by biofilms against various agar surfaces. The biofilms are first grown on gridded filter paper to ensure consistent shape across all measurements. The mature biofilm, grown for 48 hours at 23 °, is subsequently pulled by the UTM across different agar surfaces. B. The force applied by UTM is plotted against the distance the UTM crosshead traveled. The dashed line and the shaded region represent the mean and standard deviation from three different replicates, respectively. The registered force increases initially as the string stretches, but the biofilm remains stationary. Then, when the force is high enough, the biofilm begins to slide across the agar surface, at which point the force drops. C. The sliding threshold force is plotted against the agar percentage (standard error shown, n=3). We observe that the sliding threshold force increases with increasing agar percentage.

At first, the string stretches but the colony and filter do not move. However, when the applied force is large enough, the colony begins to slide across the agar surface and the applied force drops precipitously (Fig. 4B). This force threshold must be overcome for the colony to expand. We find that the value of this peak friction increases with increasing agar percentage (Fig. 4C).

### Biofilm contact angle increases with increasing friction in experiments

We next measure the contact angle of *Vibrio cholerae* WT biofilms on different agar surfaces using white light interferometry (see Methods section for more details) (16, 27–29), and plot *θ* against the corresponding value of friction measured for each agar percentage (Fig. 5A). To ensure consistency, we used the same batch of agar plates, all made following the same protocols (see Methods section for more detail), for all experiments. We find that *θ* is highly correlated with the measured friction forces (*R*^2^ = 0.88), consistent with the idea that increasing friction results in an increase in contact angle.

**Fig. 5.**
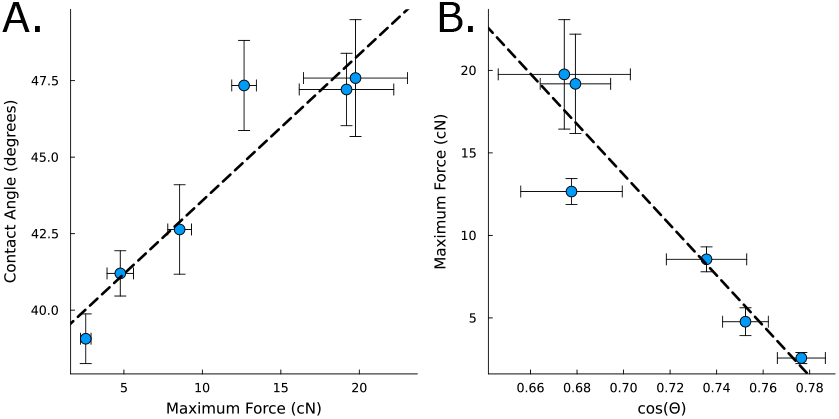
A. The contact angle, measured from experiments performed on different agar surfaces for *V. cholerae (WT)*, is plotted against the frictional force measured for the same strain. We fit a line to the means of the data and report the standard error around each point (p value of slope = 0.0057 and *R*^2^ = 0.88, n=3 and 4). B. The friction force is plotted against the cosine of the contact angle (standard error in both axes), demonstrating that the relationship discovered in our simulations holds in experiments. We fit a line to the means of the data and report the standard error around each point (p value of slope = 0.0056, *R*^2^ = 0.88)

To quantitatively test the force-balance suggested in equation 1, we now plot friction versus *cos*(*θ*) (Fig. 5B). As predicted, we observe a linearly decreasing function (*R*^2^ = 0.88), consistent with the idea that forces are dynamically balanced at the edge of the colony.

### Contact angle increases with increasing agar percentage for many bacterial species

Finally, we sought to test if these results hold beyond our WT strain of *Vibrio cholerae*. To maximize the number of different species and strains tested, we focused on measuring the contact angle as a function of agar percentage. We again grew monoclonal colonies on six different agar concentrations using four different species (*Pseudomonas aeruginosa, Aeromonas veronii, Vibrio cholerae, Bacillus subtilus*) and three strains of *V. cholerae* with varying amounts of exopolysaccharides (EPS). The three strains are EPS-, which makes no EPS, wild type (WT), which makes a moderate amount of EPS, and EPS+, which is engineered to make as much EPS as possible (30). To ensure consistency, we used the same batch of agar plates, all made following the same protocols (see Methods section for more detail), for all experiments. After 48 hours of growth, we measured the contact angle using white light interferometry.

We found that across all strains, there was a substantial increase in contact angle with increasing agar percentage (Fig. S6). The Pearson’s correlation coefficient *ρ* was ≥ 0.8 for all species and strains tested (see SI for details). These observations suggest that changes in substrate material properties strongly impact contact angle for a wide range of microbes.

## Discussion

Here, we explored the impact of friction on biofilm morphology through simulations and experiments. Simulations suggested that increasing friction results in an increased contact angle, and that contact angle is ultimately governed by a force balance reminiscent of Young’s equation. We then experimentally measured the friction between colonies and different agar percentage surfaces. Many previous works have postulated that friction increases with increasing agar percentage (e.g., (5, 9–11)); here we experimentally demonstrate that this is true. Further, we showed that contact angle increases as friction increases, and that the Young’s equation-like relation holds in experiments, too. Finally, we measured the effect of different agar surfaces on the edge geometry of biofilms across six different strains, and found that increasing agar percentage increases contact angle in all six strains. Thus, we found that friction plays an important role in determining biofilm morphology.

The idea that physical forces govern range expansion rates has important consequences (8, 25, 31–37). First, the competition between strains to capture the most surface area, and thus the most access to nutrients, is not solely decided by which strain has the faster cellular growth rate. It was previously shown that biofilm range expansion rates depend more sensitively on biofilm contact angle than on cellular doubling time (16). Here we show that friction plays a crucial role in determining contact angle, suggesting that increasing friction leads to slower range expansion rates. These results also suggest that fitness should not be defined solely to be the range expansion rate; friction slows horizontal expansion, but speeds up vertical growth. While faster horizontal growth will eventually lead to more biomass, vertical growth must be accounted for to assess the true difference in cell count, especially on short time scales.

Changing agar percentage is expected to change the bulk and surface properties of the substrate (38). Previous papers have shown that for agar percentages below 1.5%, the bulk properties substantially influence the range expansion rate via osmotic effects (4, 39, 40). Thus, we purposely worked with agar percentages ≥ 1.5% to avoid such bulk effects. Further, we previously demonstrated that single-cell doubling times do not change across agar percentages for the strains studied here (for agar percentages ≥ 1.5% see (16)).

Further, we see that increasing the amount of EPS in our *Vib-rio cholerae* strains increases the contact angle. These results appear consistent with the observation from our simulations that as cell-cell attraction increases, contact angle increases as well. In a related vein, cell adhesion to the surface is commonly measured to be larger than cell-cell adhesion (41–46), suggesting that surface forces may play a dominant role in colony development. However, direct measurement of cell-cell adhesion would be necessary to evaluate either of these concepts.

The role of mechanical forces on the propagating front of a growing colony may also have ecological and evolutionary consequences. Indeed, a slow growing strain that is able to minimize its friction with the substrate it is attached to (or cell-cell attraction) could produce a smaller contact angle and thus expand faster. Thus, physical forces, and their impact on biofilm geometry, can play as important of a role as cellular growth rates.

## Supplementary Information

### Methods

#### E. Strain preparation

We grew strains overnight in lysogeny broth (LB) media in a shaking incubator at 37 °C. Further details on modified strains of *V. cholerae* are included in Table 1. Samples grown overnight (OD1) were further diluted using PBS solution by a factor of 10^6^ - 10^7^. This allowed us to inoculate a colony starting from a single cell. The diluted sample was then inoculated on different concentration LB agar plates ranging from 1.5% to 5% w/v. All contact angle measurements were performed on colonies grown from a single cell.

**Table 1.**
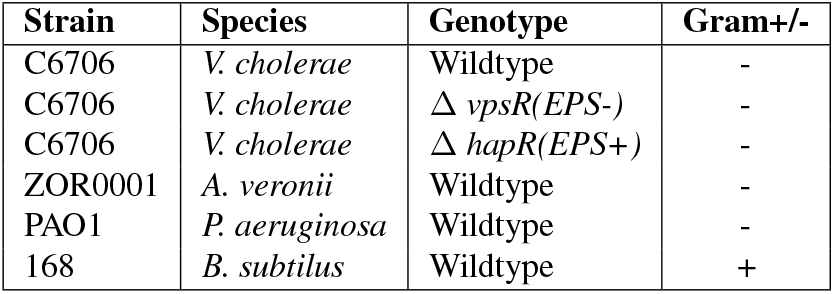
Bacterial strains used in the study.

#### F. Experimental Measurements

##### F.1. Inoculation

Inoculated colonies were incubated at 23°C for about 48 hrs for most of the experiments except for *Bacillus subtilis* which was grown for 96 hrs (due to its slower growth rate). We determined the time of growth for each strain such that we had similar sized colonies across strains, and none were wrinkled or buckled. For replicates, we selected random colonies grown on the same agar plates.

##### F.2. Image acquisition

We imaged fully grown colonies using a Zygo Zegage Pro interferometer with 50X Mirau objective. Since this objective has lateral resolution of ~174 nm/pixels, we used a 1000 by 1000 pixels field of view to image several continuous parts of the colony and then stitched these images together with Zygo’s stitching algorithm to get a full image. For all contact angle measurements, we only imaged one quarter of the colony rather than the whole colony. This reduced the amount of time for measurements and also allowed us to make measurements on all agar percentages in a single day, thus allowing a suite of experiments to be performed from agar made from the same preparation and that was the same age. The 50X Mirau objective used here has a slope limit of ~ 30 °. In cases where colonies had larger contact angles than the slope limit, we tilted the agar plates to an angle of ~ 45°to reduce data dropout and the need for interpolation in post-processing.

##### F.3. Frictional force measurement for different agar percentages

We sought to measure the macroscopic friction force exerted on a biofilm as it slides against various agar surfaces. It has been previously hypothesized that these frictional forces increase with increasing agar percentages (5, 9, 10). However, without a direct measurement of this force, more precise modeling has been difficult. Therefore, we aimed to experimentally measure this force by developing a straight-forward method using a Universal Testing Machine (UTM). We inoculated an OD1 *Vibrio cholerae wild type* onto a ME Whatman gridded filter paper. We inoculated approximately 4 µl onto the filter paper, resulting in a roughly square-shaped biofilm (due to the shape of the grids on the filter). These colonies were then left to incubate on 1.5% agar at room temperature for about 48 hours. Next, the filter paper is taken off of the agar pad it is grown on, and placed on a new agar pad with the biofilm facing downward. We then slowly moved the filter paper, along with the biofilm, across various agar surfaces (as shown in Fig. 1) using the UTM. To do so, we attached a nylon string to the Universal Testing Machine, which was equipped with a 5N load cell. The 5N load cell measures the forces up to 5N and accuracy down to 0.02 N. Initially, as the string was pulled, it stretched but the biofilm did not move. The force applied by the UTM increased until it eventually reached a threshold where the biofilm began to slide across the surface, and the force then dropped rapidly (Fig. 4B)

We determined the sliding threshold force for biofilms for six different agar concentrations. As expected, we observed that the maximum force required to start sliding across the surface increases with increasing agar percentages (Fig. 4C). Note here that we are measuring the force equivalent to static friction which is the maximum force necessary to overcome the initial resistance to sliding and induce biofilm movement across the surface.

#### G. Agent-based Model

We developed an agent-based simulation to study the growth of bacterial biofilms on a surface, focusing on how local physical forces influence global colony shape. The model was implemented in Julia using the Differential Equations package, where the integrator solved a system of 3*N* equations for *N* cells as we integrated each cell’s x position, y position, and radii. Callbacks were used to add cells and increase the size of the system (N) as division occurred. The simulation domain is a two-dimensional with a non-periodic (repulsive) boundary condition on the bottom while the left, right and top directions are unbounded.

##### Agent Properties and Initialization

Each agent *i* is characterized by its position **x**_*i*_ ∈ ℝ^2^, radius *r*_*i*_, mass *m*_*i*_, velocity **v**_*i*_, and state variables governing growth, division, and death. All model parameters are listed in Table 2. We assume an overdamped viscous environment such that momentum is not carried over time-step to time-step (see equations of motion below). At initialization, a single cell is placed on the substrate, i.e., the bottom surface.

**Table 2.** Model parameters.

| Parameter | Value | Description |
| --- | --- | --- |
| $k_c$ | 100 | Cell-cell spring constant |
| $R_{\max}$ | 3 | Maximum cell interaction distance |
| $t_{\text{final}}$ | 18 | Runtime of simulations |
| $r_{\max}$ | 0.5 | Maximum cell radius |
| $\alpha$ | 0.7 | Mean growth rate of cells |
| $A$ | [0,0.56] | Cell-cell adhesion strength |
| $L_c$ | 0.25 | Distance of max adhesion |
| $l_c$ | 0.25 | Decay length of adhesion |
| $k_w$ | 400 | Cell-substrate spring constant |
| $F$ | [0,2] | Maximum friction |
| $l_f$ | 5.0 | Decay length of friction |
| $L$ | 6.0 | Vertical nutrient limit |
| $P_f$ | 0.2 | Cell-substrate adhesion strength |
| $L_w$ | 1.0 | Distance of max cell-wall adhesion |
| $l_w$ | 0.25 | Decay length of cell-wall adhesion |
| $\gamma$ | 1 | Drag coefficient |
| $\Delta^*$ | 0.15 | Overlap threshold for growth decay |

##### Equations of Motion

At each time step, the position of each living cell is integrated according to the net force acting on it in an overdamped setting:

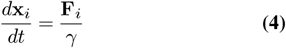

where the total force **F**_*i*_ on cell *i* is

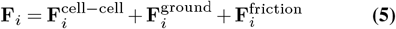

Here, 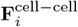 denotes the sum of all cell–cell forces, 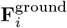 is the repulsive force from the simulation boundaries as well as the cells’ vertical adhesion to the substrate, and 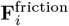 is friction with the substrate.

##### Cell–Cell Forces

Cells interact via short-range repulsion (modeling contact) and medium-range attraction (modeling adhesion). The force between two cells *i* and *j* at positions **x**_*i*_ and **x**_*j*_is

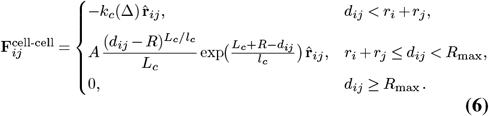

where *d*_*ij*_ = |**x**_*i*_ − **x**_*j*_| is the distance between cell centers, *R* = *r*_*i*_ + *r*_*j*_ is the sum of cell radii, so Δ = *R − d*_*ij*_ is the overlap, and *d*_*ij*_ − *R* is the distance between the outside surfaces of the two cells. 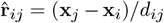 is the unit vector from *i* to *j, k* is the repulsion constant, *A* is the maximum adhesion force, and *l*_*c*_ and *L*_*c*_ control the range and decay of the attractive force. Specifically, *l*_*c*_ is the characteristic length scale over which the force exponentially drops, and *L*_*c*_ is the distance between the cell walls at which the force is maximized at *A*. Taken together, this leaves us with a force that pushes cells apart strongly when they overlap, is equilibrated when the outside surfaces are just touching, and pulls the cells together when they are slightly separated (with the adhesion strength dropping exponentially as distance increases). Once the cells are *R*_max_ apart, this is no force between them. This final condition is both for computational efficiency and because the force drops quickly with distance negating the need to calculate all pair-wise cell combinations; it is set at six maximum cell radii away. In our plots, we re-port *A*^*^ which is this maximum adhesion force divided by the average cell-cell overlapping force when cells begin to jam, i.e., 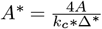 where the factor of 4 comes from the overlap being split between four neighbors (the needed number of contacts to jam symmetrical circles).

##### Cell–Wall Interaction

Cells are confined within the simulation box by a repulsive “spring” force from the boundaries:

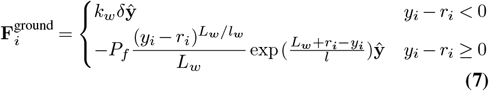

Where *y*_*i*_ is the vertical distance from a cell’s center to the ground, *δ* is the overlap of cell *i* with the wall (*δ* = **x**_*i*_ · *ŷ* − *r*_*i*_), *k*_*w*_ is the wall repulsion constant. When cells do not overlap with the wall, they experience a small attractive force pulling downward of the same form as cell-cell adhesion, but the equilibrium position is when cells rest on the ground. This force is maximized at 1.0 microns away from this equlibrium and then likewise decays exponentially. The maximum of this force is held constant across simulations.

##### Friction with the Substrate

We consider friction between cells and the substrate; it is strongest when the cells are near the substrate but decays exponentially with length *l*_*f*_. Friction opposes the net lateral (horizontal) force, but is capped at a maximum value *F*. Thus, the friction force on cell *i* is given by:

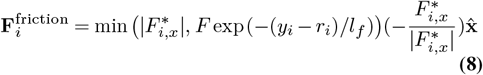

Here, 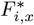 is the total force, outside of friction, in the *x*-direction on cell 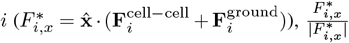 is the the current direction of the lateral force, |**F**_*i,x*_| is its magnitude, and *F* is the maximum friction force set by the substrate. We note that the ground force only has vertical components, so the friction is then only affected by cell-cell interactions. Friction does not affect vertical forces. In our plots, we report *F* ^*^ which is this maximum friction force divided by the average cell-cell overlapping force when cells begin to jam ie 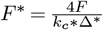 where the factor of 4 comes from the overlap being split between eight neighbors and then we half the overlap as this is ‘just before’ jamming.

##### Growth, Division, and Death

The cells’ area grow exponentially at rate *α* when in proximity to the substrate and thus the nutrients (*y < L*)(27), up to a maximal area *A*_max_. Further, each cell is given a random growth rate uniformly distributed around 0.7 with a 10% spread to avoid synchronous division. Note, this means that the global growth rate is lower than 0.7 as there is a non-linear relation between a spread in the individual growth rates and the emergent population level growth. Each cell also has a slower growth rate when the sum of the overlapping cells exceeds a threshold Δ^*^, which we set at 0.15 microns.

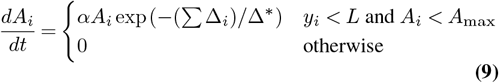

Once a cell reaches its maximum area, it divides into two daughter cells with the same area, offset by a distance such that their outer membranes are just touching. Further, the two daughter cells’ centers make a line parallel to the ground.

##### Simulation Protocol

At each simulation step: (1) pairwise forces are computed using a cell list for efficiency (i.e., in a cell list, cell positions are binned into groups, such that, for example, all of the cells with centers between 0 *< x <* 1 and 0 *< y <* 1 are stored together; thus, when computing pairwise forces we only search for neighbors within the nearby bins rather than among all cells, and we chose bin sizes such that we do not miss any cells within the maximum interaction range *R*_max_); (2) forces are summed and used to update the velocities which are passed into an integrator to update positions (Julia’s Tsit5 integrator which is an explicit, adaptive Runge–Kutta integrator of order 5 with an embedded order-4 error estimator); (3) growth of each cell; radii are passed into the integrator to increase them; and (4) division occurs if a cell’s radius reaches the maximum value. When cell divides, the system is resized to hold one more cell (three equations are produced: one for the x position, one for the y position, and one for the radius). This modeling framework enables us to systematically explore the impact of substrate friction, cell–cell adhesion, and other physical parameters on emergent biofilm morphology, particularly the contact angle and range expansion rate.

**Fig. S6.**
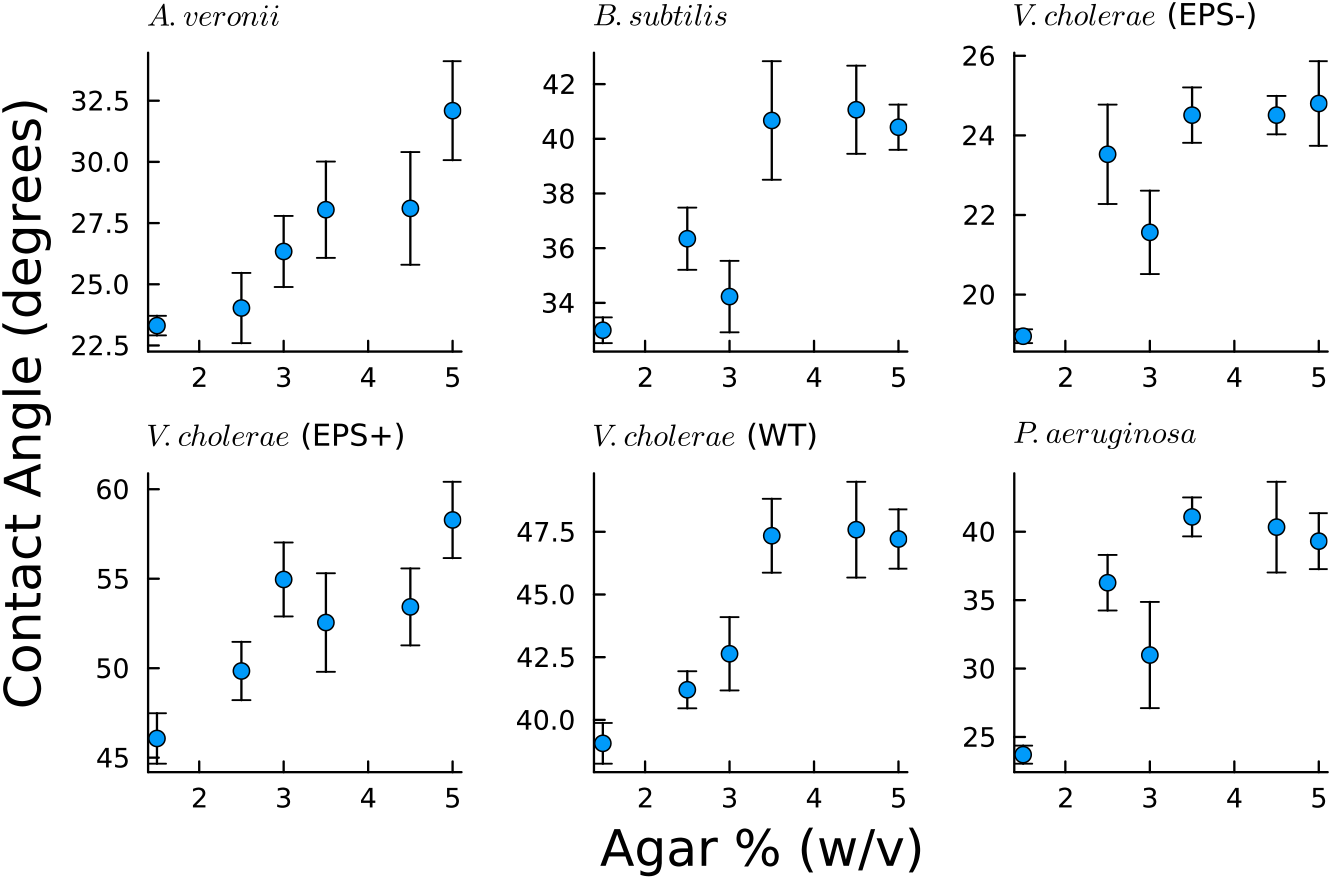
Contact angle is plotted against agar percentage for four species and three strains of *Vibrio cholerae*. We observe that as the agar percentage increases, contact angle consistently also increases as well. The species, and in some cases, strain, is indicated at the top of each panel. All y-axes are contact angle in degrees; all x-axes are Agar Percentage.

**Fig. S7.**
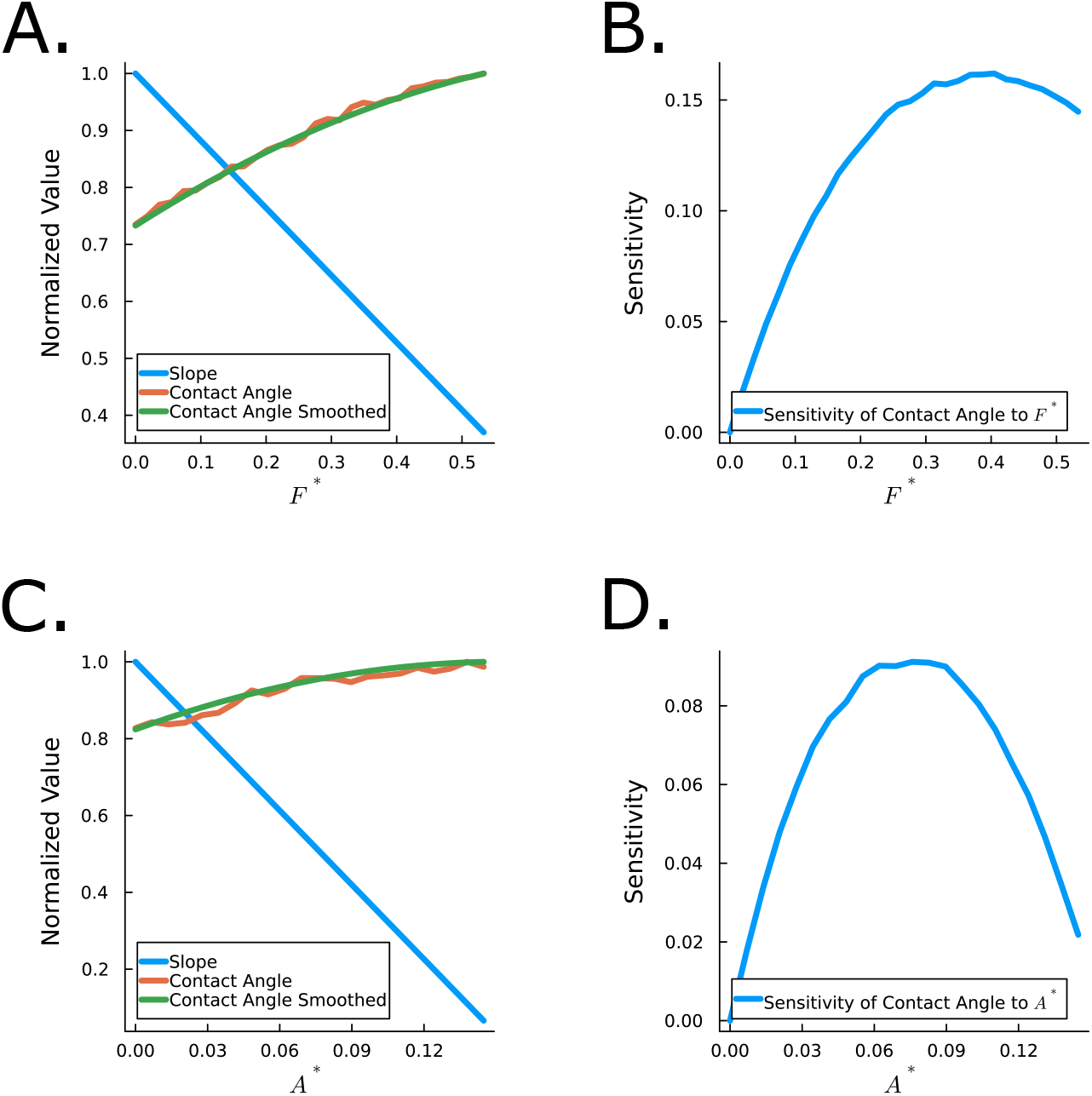
A. The same data used in figure 2 where we have added a smoothed line using a window moving degree 2 polynomial to smooth and find the derivative of contact angle with respect to friction and attraction B. The sensitivity of the contact angle with respect to friction and attraction, we see that friction impact the contact angle more for all forces simulated here.

**Fig. S8.**
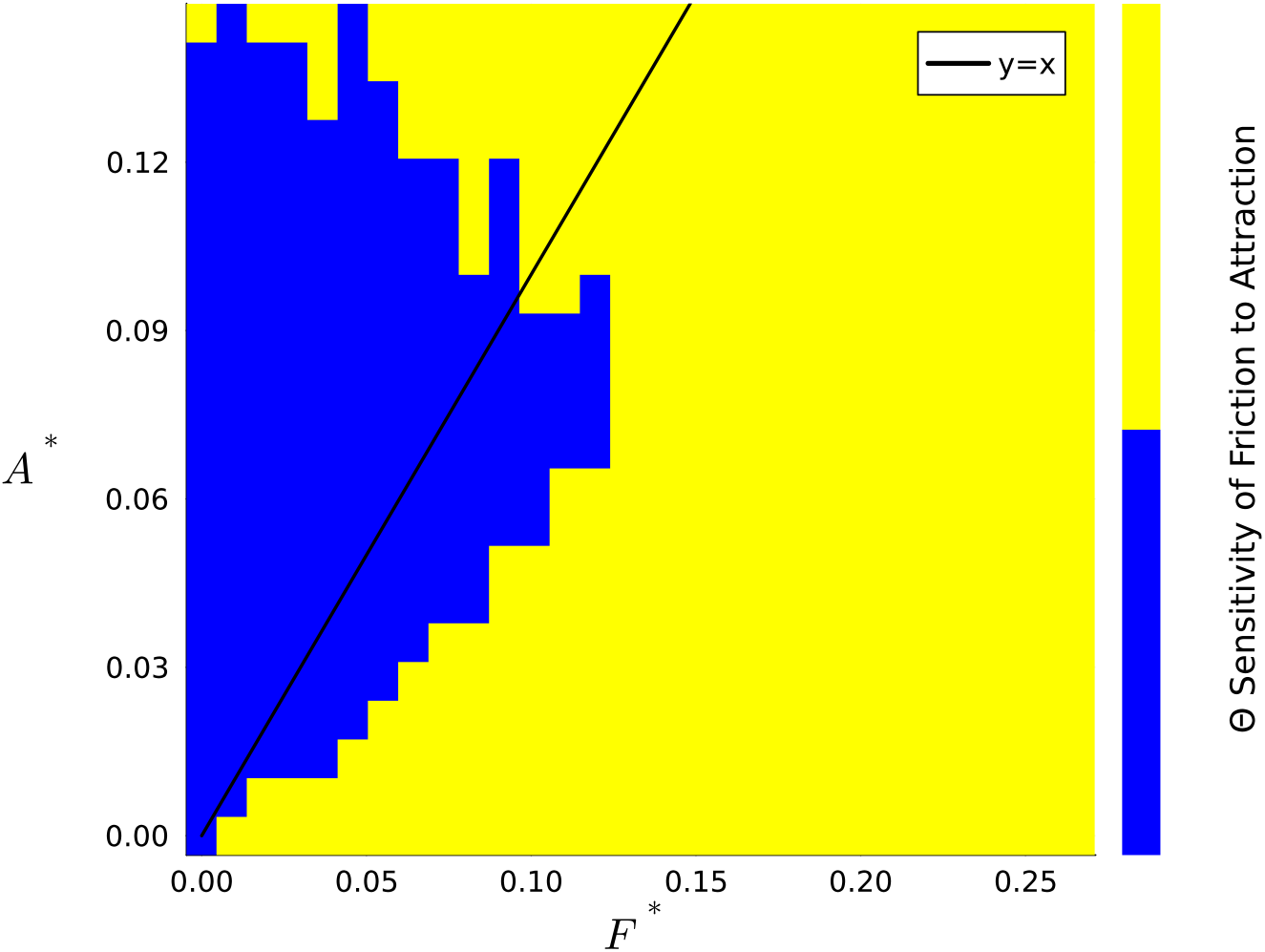
We measured the sensitivity of the contact angle as influenced by friction and cell-cell attraction (as shown in S7)) across a variety of attraction and friction forces. Here we plot a binarized image of the contact angle’s sensitivity of friction to attraction where yellow implies friction is more sensitive and blue attraction. We see that 72% of the tested values were more sensitive to friction than attraction

## Bibliography

1. Hans-Curt Flemming and Stefan Wuertz. Bacteria and archaea on Earth and their abundance in biofilms. Nature Reviews Microbiology, 17(4):247–260, 2019. Publisher: Nature Publishing Group.

2. Ailing Zhao, Jiazheng Sun, and Yipin Liu. Understanding bacterial biofilms: From definition to treatment strategies. Frontiers in Cellular and Infection Microbiology, 13:1137947, April 2023. ISSN 2235-2988. doi: 10.3389/fcimb.2023.1137947.

3. Simon R. Law, Falko Mathes, Amy M. Paten, Pamela A. Alexandre, Roshan Regmi, Cameron Reid, Azadeh Safarchi, Shaktivesh Shaktivesh, Yanan Wang, Annaleise Wilson, Scott A. Rice, and Vadakattu V. S. R. Gupta. Life at the borderlands: microbiomes of interfaces critical to One Health. FEMS microbiology reviews, 48(2):fuae008, March 2024. ISSN 1574-6976. doi: 10.1093/femsre/fuae008.

4. Merrill E Asp, Minh-Tri Ho Thanh, Danielle A Germann, Robert J Carroll, Alana Franceski, Roy D Welch, Arvind Gopinath, and Alison E Patteson. Spreading rates of bacterial colonies depend on substrate stiffness and permeability. PNAS nexus, 1(1):pgac025, 2022. Publisher: Oxford University Press.

5. Mya R Warren, Hui Sun, Yue Yan, Jonas Cremer, Bo Li, and Terence Hwa. Spatiotemporal establishment of dense bacterial colonies growing on hard agar. Elife, 8:e41093, 2019. Publisher: eLife Sciences Publications Limited.

6. Agnese Seminara, Thomas E Angelini, James N Wilking, Hera Vlamakis, Senan Ebrahim, Roberto Kolter, David A Weitz, and Michael P Brenner. Osmotic spreading of Bacillus subtilis biofilms driven by an extracellular matrix. Proceedings of the National Academy of Sciences, 109(4):1116–1121, 2012. Publisher: National Acad Sciences.

7. Baochi Nguyen, Arpita Upadhyaya, Alexander van Oudenaarden, and Michael P Brenner. Elastic instability in growing yeast colonies. Biophysical journal, 86(5):2740–2747, 2004. Publisher: Elsevier.

8. Chenyi Fei, Sheng Mao, Jing Yan, Ricard Alert, Howard A Stone, Bonnie L Bassler, Ned S Wingreen, and Andrej Košmrlj. Nonuniform growth and surface friction determine bacterial biofilm morphology on soft substrates. Proceedings of the National Academy of Sciences, 117(14):7622–7632, 2020. Publisher: National Acad Sciences.

9. Yusuf Ilker Yaman, Esin Demir, Roman Vetter, and Askin Kocabas. Emergence of active nematics in chaining bacterial biofilms. Nature communications, 10(1):2285, 2019. Publisher: Nature Publishing Group UK London.

10. Japinder Nijjer, Changhao Li, Mrityunjay Kothari, Thomas Henzel, Qiuting Zhang, Jung-Shen B Tai, Shuang Zhou, Tal Cohen, Sulin Zhang, and Jing Yan. Biofilms as self-shaping growing nematics. Nature Physics, pages 1–9, 2023. Publisher: Nature Publishing Group UK London.

11. Matthew AA Grant, Bartlomiej Waclaw, Rosalind J Allen, and Pietro Cicuta. The role of mechanical forces in the planar-to-bulk transition in growing Escherichia coli microcolonies. Journal of The Royal Society Interface, 11(97):20140400, 2014. Publisher: The Royal Society.

12. He Xu, Anne E Murdaugh, Wei Chen, Katherine E Aidala, Megan A Ferguson, Eileen M Spain, and Megan E Núñez. Characterizing pilus-mediated adhesion of biofilm-forming E. coli to chemically diverse surfaces using atomic force microscopy. Langmuir, 29(9):3000– 3011, 2013. Publisher: ACS Publications.

13. Xavier Trepat Guixer and Erik Sahai. Mesoscale physical principles of collective cell organization. Nature Physics, 2018, vol. 14, p. 671–682, 2018. Publisher: Nature Publishing Group.

14. M Wang, D Xu, K Ravi-Chandar, and KM Liechti. On the development of a mesoscale friction tester. Experimental mechanics, 47:123–131, 2007. Publisher: Springer.

15. Xi Shi and Andreas A Polycarpou. Measurement and modeling of normal contact stiffness and contact damping at the meso scale. J. Vib. Acoust., 127(1):52–60, 2005.

16. Aawaz R Pokhrel, Gabi Steinbach, Adam Krueger, Thomas C Day, Julianne Tijani, Brian K Hammer, and Peter J Yunker. The biophysical basis of bacterial colony growth. bioRxiv, pages 2023–11, 2023. Publisher: Cold Spring Harbor Laboratory.

17. Thomas Young. Iii. An essay on the cohesion of fluids. Philosophical Transactions of the Royal Society of London, 95:65–87, January 1805. doi: 10.1098/rstl.1805.0005. Publisher: Royal Society.

18. Jaroslaw W Drelich, Ludmila Boinovich, Emil Chibowski, Claudio Della Volpe, Lucyna Holysz, Abraham Marmur, and Stefano Siboni. Contact angles: history of over 200 years of open questions. Surface Innovations, 8(1-2):3–27, February 2020. ISSN 2050-6252. doi: 10.1680/jsuin.19.00007. Publisher: ICE Publishing.

19. Raimo Hartmann, Praveen K Singh, Philip Pearce, Rachel Mok, Boya Song, Francisco Díaz-Pascual, Jörn Dunkel, and Knut Drescher. Emergence of three-dimensional order and structure in growing biofilms. Nature physics, 15(3):251–256, 2019. Publisher: Nature Publishing Group UK London.

20. Zhaowei Jiang, Thomas Nero, Sampriti Mukherjee, Rich Olson, and Jing Yan. Searching for the Secret of Stickiness: How Biofilms Adhere to Surfaces. Frontiers in Microbiology, 12, July 2021. ISSN 1664-302X. doi: 10.3389/fmicb.2021.686793. Publisher: Frontiers.

21. Andrea J. Liu and Sidney R. Nagel. The Jamming Transition and the Marginally Jammed Solid. Annual Review of Condensed Matter Physics, 1(Volume 1, 2010):347–369, August 2010. ISSN 1947-5454, 1947-5462. doi: 10.1146/annurev-conmatphys-070909-104045. Publisher: Annual Reviews.

22. Marie-Cécilia Duvernoy, Thierry Mora, Maxime Ardré, Vincent Croquette, David Bensimon, Catherine Quilliet, Jean-Marc Ghigo, Martial Balland, Christophe Beloin, Sigolène Lecuyer, and Nicolas Desprat. Asymmetric adhesion of rod-shaped bacteria controls microcolony morphogenesis. Nature Communications, 9(1):1120, March 2018. ISSN 2041-1723. doi: 10.1038/s41467-018-03446-y. Publisher: Nature Publishing Group.

23. Denis Boyer, William Mather, Octavio Mondragón-Palomino, Sirio Orozco-Fuentes, Tal Danino, Jeff Hasty, and Lev S Tsimring. Buckling instability in ordered bacterial colonies. Physical Biology, 8(2):026008, March 2011. ISSN 1478-3975. doi: 10.1088/1478-3975/8/2/026008.

24. Farzan Beroz, Jing Yan, Yigal Meir, Benedikt Sabass, Howard A Stone, Bonnie L Bassler, and Ned S Wingreen. Verticalization of bacterial biofilms. Nature physics, 14(9):954–960, 2018. Publisher: Nature Publishing Group.

25. Alice Cont, Tamara Rossy, Zainebe Al-Mayyah, and Alexandre Persat. Biofilms deform soft surfaces and disrupt epithelia. Elife, 9:e56533, 2020. Publisher: eLife Sciences Publications, Ltd.

26. Edda Klipp, Wolfram Liebermeister, Christoph Wierling, Axel Kowald, Hans Lehrach, and Ralf Herwig. Systems Biology. Wiley-VCH Verlag GmbH & Co. KGaA, Weinheim, 2009.

27. Pablo Bravo, Siu Lung Ng, Kathryn A MacGillivray, Brian K Hammer, and Peter J Yunker. Vertical growth dynamics of biofilms. Proceedings of the National Academy of Sciences, 120(11):e2214211120, 2023. Publisher: National Acad Sciences.

28. Pablo J Bravo and Peter J Yunker. Fluctuations and freezing of biofilm-air interfaces. bioRxiv, pages 2024–05, 2024. Publisher: Cold Spring Harbor Laboratory.

29. Arben Kalziqi, David Yanni, Jacob Thomas, Siu Lung Ng, Skanda Vivek, Brian K. Hammer, and Peter J. Yunker. Immotile Active Matter: Activity from Death and Reproduction. Physical Review Letters, 120(1):018101, January 2018. doi: 10.1103/PhysRevLett.120.018101. Publisher: American Physical Society.

30. Brian K. Hammer and Bonnie L. Bassler. Quorum sensing controls biofilm formation in Vibrio cholerae. Molecular Microbiology, 50(1):101–104, 2003. ISSN 1365-2958. doi: 10.1046/j.1365-2958.2003.03688.x. _eprint: https://onlinelibrary.wiley.com/doi/pdf/10.1046/j.13652958.2003.03688.x.

31. Jakub A. Kochanowski, Bobby Carroll, Merrill E. Asp, Emma C. Kaputa, and Alison E. Patteson. Bacteria Colonies Modify Their Shear and Compressive Mechanical Properties in Response to Different Growth Substrates. ACS Applied Bio Materials, 7(12):7809–7817, December 2024. doi: 10.1021/acsabm.3c00907. Publisher: American Chemical Society.

32. Jing Yan, Chenyi Fei, Sheng Mao, Alexis Moreau, Ned S Wingreen, Andrej Košmrlj, Howard A Stone, and Bonnie L Bassler. Mechanical instability and interfacial energy drive biofilm morphogenesis. Elife, 8:e43920, 2019. Publisher: eLife Sciences Publications, Ltd.

33. Qiuting Zhang, Jian Li, Japinder Nijjer, Haoran Lu, Mrityunjay Kothari, Ricard Alert, Tal Cohen, and Jing Yan. Morphogenesis and cell ordering in confined bacterial biofilms. Proceedings of the National Academy of Sciences, 118(31):e2107107118, 2021. Publisher: National Acad Sciences.

34. Alexandre Persat, Carey D. Nadell, Minyoung Kevin Kim, Francois Ingremeau, Albert Siryaporn, Knut Drescher, Ned S. Wingreen, Bonnie L. Bassler, Zemer Gitai, and Howard A. Stone. The Mechanical World of Bacteria. Cell, 161(5):988–997, May 2015. ISSN 0092-8674, 1097-4172. doi: 10.1016/j.cell.2015.05.005. Publisher: Elsevier.

35. Oskar Hallatschek, Sujit S. Datta, Knut Drescher, Jörn Dunkel, Jens Elgeti, Bartek Waclaw, and Ned S. Wingreen. Proliferating active matter. Nature Reviews Physics, 5(7):407–419, July 2023. ISSN 2522-5820. doi: 10.1038/s42254-023-00593-0. Publisher: Nature Publishing Group.

36. Alejandro Martínez-Calvo, Tapomoy Bhattacharjee, R Kőnane Bay, Hao Nghi Luu, Anna M Hancock, Ned S Wingreen, and Sujit S Datta. Morphological instability and roughening of growing 3D bacterial colonies. Proceedings of the National Academy of Sciences, 119(43): e2208019119, 2022. Publisher: National Acad Sciences.

37. Steffen Geisel, Eleonora Secchi, and Jan Vermant. The role of surface adhesion on the macroscopic wrinkling of biofilms. eLife, 11:e76027, June 2022. ISSN 2050-084X. doi: 10.7554/eLife.76027. Publisher: eLife Sciences Publications, Ltd.

38. Valéry Normand, Didier L Lootens, Eleonora Amici, Kevin P Plucknett, and Pierre Aymard. New insight into agarose gel mechanical properties. Biomacromolecules, 1(4):730–738, 2000. Publisher: ACS Publications.

39. Jing Yan, Carey D. Nadell, Howard A. Stone, Ned S. Wingreen, and Bonnie L. Bassler. Extracellular-matrix-mediated osmotic pressure drives Vibrio cholerae biofilm expansion and cheater exclusion. Nature Communications, 8(1):327, August 2017. ISSN 2041-1723. doi: 10.1038/s41467-017-00401-1.

40. Alison E. Patteson, Merrill E. Asp, and Paul A. Janmey. Materials science and mechanosensitivity of living matter. Applied Physics Reviews, 9(1):011320, March 2022. ISSN 1931-9401. doi: 10.1063/5.0071648.

41. Tal Duanis-Assaf and Meital Reches. Factors influencing initial bacterial adhesion to antifouling surfaces studied by single-cell force spectroscopy. iScience, 27(2):108803, January 2024. ISSN 2589-0042. doi: 10.1016/j.isci.2024.108803.

42. M.J. Chen, Z. Zhang, and T.R. Bott. Direct measurement of the adhesive strength of biofilms in pipes by micromanipulation. Biotechnology Techniques, 12(12):875–880, December 1998. ISSN 1573-6784. doi: 10.1023/A:1008805326385.

43. René Riedel Garima Rani, and Anupam Sengupta. Bacterial Adhesion on Soft Surfaces: The Dual Role of Substrate Stiffness and Bacterial Growth Stage. Microorganisms, 13(3): 637, March 2025. ISSN 2076-2607. doi: 10.3390/microorganisms13030637.

44. Xing () Jin and Jeffrey S. Marshall. Influence of cell interaction forces on growth of bacterial biofilms. Physics of Fluids, 32(9):091902, September 2020. ISSN 1070-6631. doi: 10.1063/5.0021126.

45. Cécile Formosa-Dague, Pietro Speziale, Timothy J. Foster, Joan A. Geoghegan, and Yves F. Dufrêne. Zinc-dependent mechanical properties of Staphylococcus aureus biofilm-forming surface protein SasG. Proceedings of the National Academy of Sciences, 113(2):410–415, January 2016. doi: 10.1073/pnas.1519265113. Publisher: Proceedings of the National Academy of Sciences.

46. Paula Parreira and M. Cristina L. Martins. The biophysics of bacterial infections: Adhesion events in the light of force spectroscopy. The Cell Surface, 7:100048, December 2021. ISSN 2468-2330. doi: 10.1016/j.tcsw.2021.100048.

